# Unraveling the Molecular and Physiological Roles of Signal Peptide Peptidase A in *Flavobacterium columnare*

**DOI:** 10.1101/2025.09.26.678728

**Authors:** Ruoxi Zhu, Liang Zhong, Yuying Xun, Shucheng Zheng, Yongtao Zhu, Wenlong Cai

**Affiliations:** Department of Infectious Disease and Public Health, Jockey Club College of Veterinary Medicine and Life Science, City University of Hong Kong, Kowloon Tong, Hong Kong SAR, China; Department of Biosciences and Bioinformatics, School of Science, Xi’an Jiaotong-Liverpool University, Suzhou, Jiangsu, China; Wisdom Lake Academy of Pharmacy, Xi’an Jiaotong-Liverpool University, Suzhou Jiangsu, China; Key Laboratory of Fishery Drug Development, Ministry of Agriculture and Rural Affairs, Guangdong Provincial Key Laboratory of Aquatic Animal Immunology and Sustainable Aquaculture, Pearl River Fisheries Research Institute, Chinese Academy of Fishery Sciences, Guangzhou, 510380, China

**Keywords:** Bacterial virulence, SppA, outer membrane vesicle, columnaris disease, aquatic pathogen, sustainable aquaculture, fish health

## Abstract

Columnaris disease, caused by *Flavobacterium columnare*, represents one of the most economically devastating bacterial infections in global freshwater aquaculture. Despite its significant impact, the molecular mechanisms underlying *F. columnare* pathogenesis remain largely unexplored due to the challenge in targeted genomic manipulation. Signal peptide peptidase A (SppA) plays a crucial role in bacterial protein secretion by degrading residual signal peptides after protein translocation, yet its function in *F. columnare* physiology and virulence has not been characterized. Here, we employed a targeted gene deletion approach to investigate the role of *sppA* in *F. columnare*. The Δ*sppA* mutant exhibited pleiotropic phenotypes including increased outer membrane vesicle (OMV) production (3.8-fold higher compared to the wild type), reduced biofilm formation, and loss of gliding motility. Transcriptomic analysis of the Δ*sppA* mutant revealed significant upregulation of genes involved in membrane stress response and efflux pump system, including *algU*, *osmC* and the genes in the MacAB-TolC efflux system, compared to the wild-type state. Importantly, the artificial infection experiment demonstrated the mutant’s significantly attenuated virulence in freshwater Medaka (*Oryzias latipes*), with a 20% higher survival rate of fish compared to the wild type. Our findings reveal that SppA is essential for maintaining membrane homeostasis in *F. columnare* and serves as one of the virulence factors during columnaris infection. These results provide important insights into the biological function of the *sppA* gene in *F. columnare* and highlight the complex relationship between bacterial protein secretion, membrane integrity, and pathogenesis.

**Importance:** *F. columnare* causes significant economic loss in freshwater aquaculture. Understanding the molecular mechanisms underlying *F. columnare* pathogenesis is crucial for developing new ways for disease control. Our findings reveal that SppA is essential for gliding motility, adhesion, biofilm formation and maintaining membrane homeostasis in *F. columnare,* which serves as one of the virulence factors during columnaris infection. In addition, outer membrane vesicles (OMVs) and MacA/MacB/TolC tripartite efflux pump served as a compensatory mechanism for enhanced peptide metabolites secretion to manage the accumulation of misfolded proteins resulting from the sppA deficiency. These results provide important insights into the biological function of the sppA gene in *F. columnare* and highlight the complex relationship between bacterial protein secretion, membrane integrity, and pathogenesis.

## Introduction

*Flavobacterium columnare* is a Gram-negative, rod-shaped bacterium that causes columnaris disease in freshwater fish worldwide [1, 2]. This disease affects diverse fish species, including catfish, tilapia, and ornamental fish, resulting in substantial economic losses to aquaculture industries [3, 4]. The pathogen is characterized by its ability to form biofilms, exhibit gliding motility, and secrete various extracellular proteins through the type IX secretion system (T9SS) [5–7]. Despite its economic importance, the molecular mechanisms of *F. columnare* pathogenesis remain poorly understood due to its challenge in targeted genomic manipulation [8], and effective control strategies are limited due to the incomplete characterization of virulence factors.

Protein secretion is a key determinant of bacterial pathogenicity [9]. The secretion of virulence factors involves the coordinated action of multiple systems. The Sec system serves as the basic transport machinery, responsible for transporting newly synthesized proteins across the inner membrane into the periplasm [10]. The T9SS is a secretion system unique to the *Bacteroidota* phylum, responsible for transporting various proteins including virulence factors to the cell surface or to the extracellular environment [11, 12]. Although the T9SS selects substrates by recognizing C-terminal domain (CTD) signals, these substrates must first be transported to the periplasm via the Sec system [13]. Therefore, the function of the T9SS depends on the normal operation of the Sec system. The continuous flow of proteins through the Sec system generates signal peptide fragments embedded in the membrane, which must be effectively cleared to prevent blockage of the transport channel and maintain secretory capacity.

Signal peptide peptidase A (SppA) is a membrane-bound protease that plays a crucial role in the processing of secreted proteins. These proteins are initially produced with an N-terminal signal peptide that guides their targets to cross the cytoplasmic membrane. Following protein translocation via the Sec system, signal peptidase cleaves the signal peptide, leaving it embedded in the membrane. SppA then performs intramembrane proteolysis, cleaving within the hydrophobic core of these residual signal peptides. SppA is thus responsible for degrading these residual signal peptides and preventing their accumulation, which could otherwise disrupt membrane function and interfere with continued protein translocation [14]. Studies in *Bacillus subtilis* have demonstrated that *sppA* deletion leads to signal peptide accumulation, resulting in membrane stress, impaired Sec translocon function, and reduced protein secretion efficiency [15]. Given that T9SS-dependent virulence factors must first traverse the inner membrane via the Sec system, SppA function is potentially critical for maintaining the protein secretion capacity necessary for *F. columnare* pathogenesis.

In a previous study, we developed an attenuated vaccine strain of *F. columnare* through antibiotic-induced mutagenesis and identified missense mutations in the *F. columnare sppA* gene [16]. Another objective of this study is to investigate whether the *sppA* mutation directly contributed to the loss of virulence or was merely an incidental change. Given the essential role of protein secretion in *F. columnare* virulence and the dependence of T9SS substrates on Sec-mediated translocation, we hypothesized that SppA is a potential virulence factor for *F. columnare* pathogenesis by ensuring efficient clearance of signal peptides and maintaining membrane homeostasis. Therefore, this study aimed to investigate the biological functions of SppA in *F. columnare* and determine its contribution to bacterial virulence through the construction and characterization of a *sppA* deletion mutant.

## Results

### Generation of the *sppA* deletion mutant and the complemented strain

The suicide plasmid pMS75 was successfully constructed and employed to create the intragenic deletion mutant Δ*sppA* through double-crossover homologous recombination. Furthermore, the full-length *sppA* gene, along with its native promoter region, was cloned into the shuttle vector pCP23 and introduced into the Δ*sppA* mutant via conjugation, resulting in the complementation strain C-*sppA*. The successful construction of the *sppA* deletion mutant and the complementation strain was verified using PCR and qRT-PCR analysis (Fig. 1).

**Figure 1.**
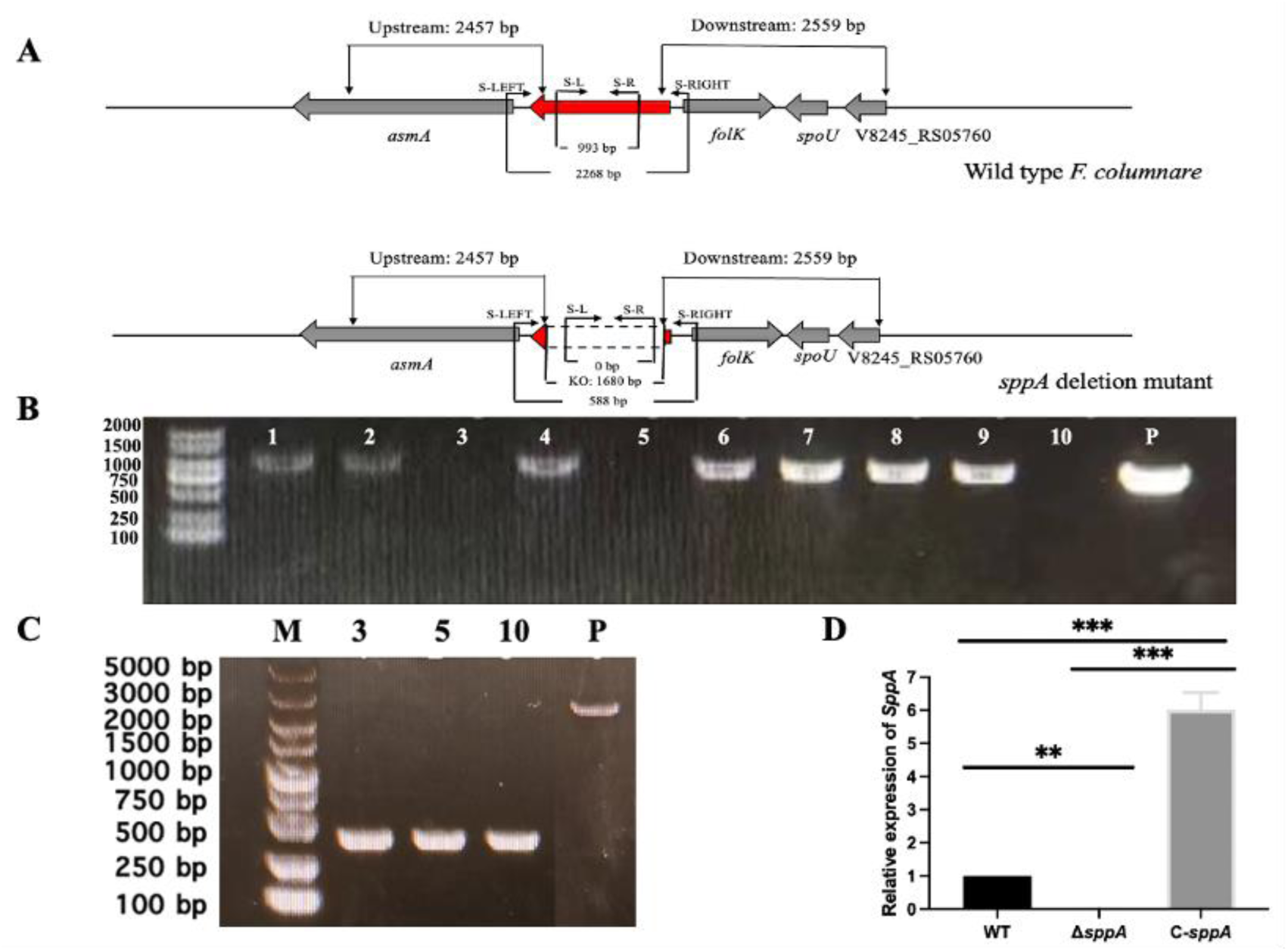
Construction of the *sppA* mutant and the complemented strain. **(A)** Genomic map of *F. columnare sppA* and its deletion. The numbers shown above and below the map indicate the length of the amplified sequence. Primer binding sites used to generate deletion or complementary constructs in PCR reactions are shown above the map. Deleted regions in the mutant are indicated by dashed lines. **(B)** PCR screening of the mutant strains (Δ*sppA*). The primers S-L and S-R, which were designed within the deleted region of *sppA*, did not amplify a product from the Δ*sppA* deletion mutant, while the wild-type strain amplified the corresponding bands (993 bp). Lanes 1, 2, 4, 6, 7, 8, and 9 correspond to colonies that produced the expected wild-type band, indicating the presence of the wild-type allele. Lanes 3, 5, and 10 did not yield a PCR product, which is consistent with the Δ*sppA* deletion genotype, but may also result from PCR failure; therefore, additional confirmation (see panel C) was performed. Lane “P” contains wild-type genomic DNA as the PCR template, serving as a wild-type control. **(C)** PCR confirmation of *sppA* gene deletion mutants. The forward primer S-LEFT and reverse primer S-RIGHT, designed within the upstream and downstream sequences of the deleted region, were used in the PCR. The bands amplified in the deletion mutants were significantly smaller than those obtained in the wild-type strain, with the difference being the number of DNA bases in the deleted region (1680 bp). Lanes 3, 5, 10 correspond to three independent Δ*sppA* mutants identified in the PCR screening shown in panel B, while lane “P” contains wild-type genomic DNA as the PCR template. The two PCR tests indicated that these strains (3, 5, and 10) were the correct *sppA* gene deletion mutants (Δ*sppA*). **(D)** Detection of the *sppA* expression. The expression levels of the *sppA* gene in the wild-type *F. columnare*, Δ*sppA,* and C-*sppA*, were determined by qRT-PCR. The data are presented as mean ± standard deviation (SD) from three independent biological replicates. ** *P < 0.01*, *** *P < 0.001*.

### The *sppA* mutant exhibited reduced growth rate under standard culture conditions

Wild-type *F. columnare* and the *sppA* mutant were grown in MS medium, and biomass accumulation was tracked as OD_600_. The strains showed comparable lag phases (*P* > 0.05). Thereafter, the curves diverged: the wild type reached significantly higher OD_600_ than Δ*sppA* at 27 and 36 h (log phase) and at 45 and 54 h (stationary phase), with C-*sppA* partially restoring the wild-type phenotype. The mutant reached the stationary phase faster. These pattern points to a role of *sppA* in bacterial fitness under nutrient limitation upon reaching the stationary phase (Fig. 2).

**Figure 2.**
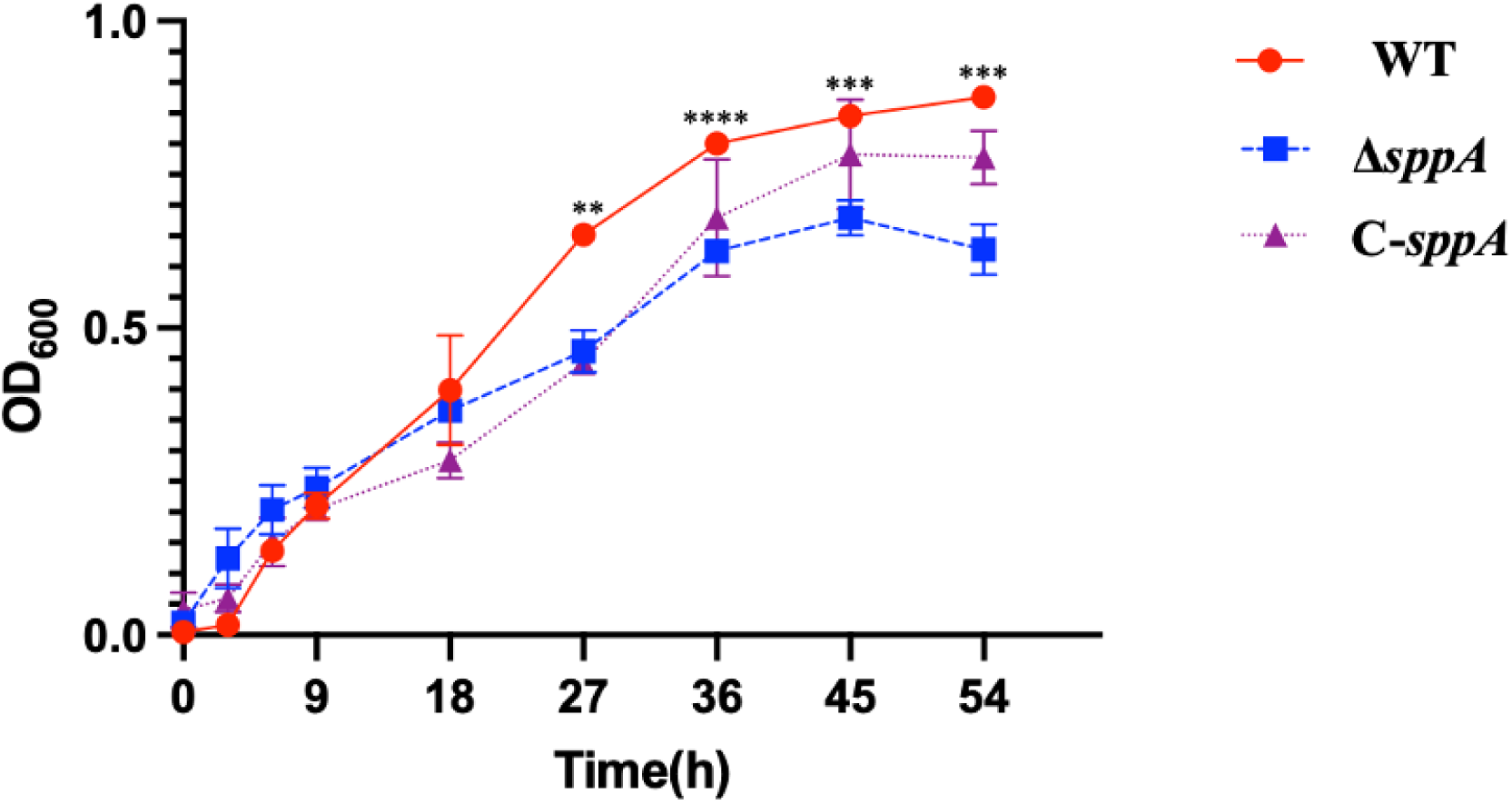
Growth curves of wild-type *F. columnare* (WT, red), Δ*sppA* mutant (Δ*sppA,* blue), and complemented strain (C-*sppA,* purple), respectively. The bacteria were cultured in MS broth at 28°C with shaking (125 rpm), and OD_600_ was measured every 9 h for 54 h. Data are presented as means ± SD. ** *P < 0.01*, *** *P < 0.001*, or **** *P < 0.0001* represents a highly significant difference, extremely significant difference, or most extremely significant difference between the wild-type *F. columnare* and the Δ*sppA* mutant.

### Deletion of *sppA* does not alter antibiotic resistance in *F. columnare*

We investigated the effect of the *sppA* gene deletion on bacterial antibiotic resistance. Here, we detected changes in the MIC of commonly used antibiotics for the treatment of *F. columnare* infections in aquaculture. However, the results showed that the deletion of *sppA* gene did not alter bacterial resistance to commonly used antibiotics (i.e. OTC, ENRO, FF) (Table S1).

### Effect of deletion of the *sppA* gene on gliding motility and colony spreading

Wild-type cells glide across glass surfaces, whereas the Δ*sppA* mutant displays defective gliding motility on glass, and complementation of the Δ*sppA* mutant restores this motility (Fig. 3A and Fig. 3B and Supplementary Movie S1). Due to gliding movements, the cells of the wild-type and complemented strains developed thin, spreading colonies on MS agar, while the cells of the Δ*sppA* mutant produced non-spreading colonies (Fig. 4).

**Figure 3.**
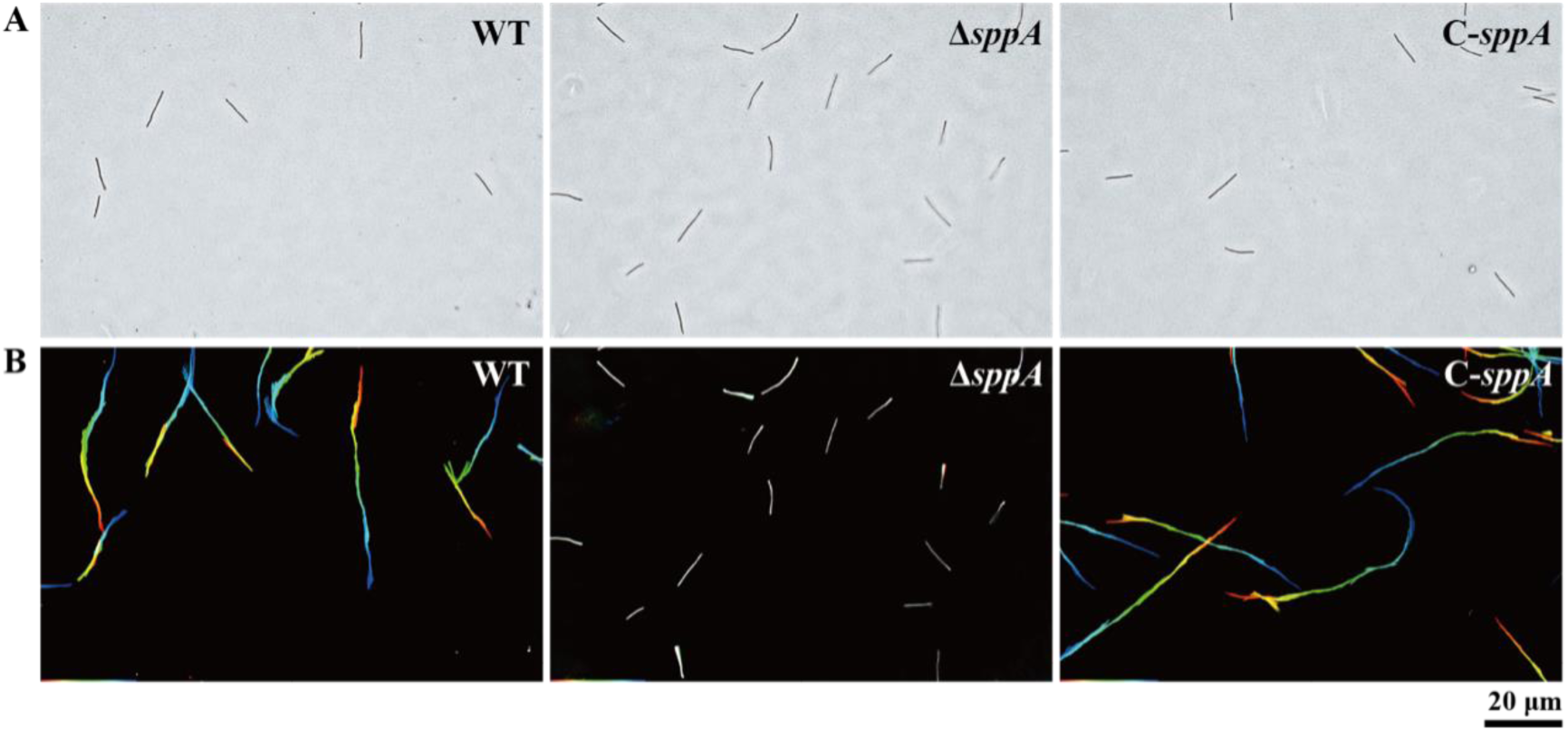
Gliding motility of wild-type *F. columnare*, Δ*sppA* mutant, and complemented strain on glass surfaces. Wild-type *F. columnare*, the Δ*sppA* mutant, and the Δ*sppA* strain carrying wild-type *sppA* on pCP23 (C-*sppA*) were cultured in 1/10 MS medium at 28°C overnight with shaking; 10 μl of each culture was added to glass tunnel slides, and cell motility was observed using a Nikon Ci-L plus microscope, with single-frame images colored from red (time zero) through orange, yellow, green, cyan, to blue (30 seconds) and integrated to display the ‘rainbow trace’ of gliding cells. **(A)** The initial frame of the motility video for each strain. **(B)** The corresponding 30-second rainbow trajectories, where white cells indicate minimal movement, the 20 μm scale bar applies to all images, and the rainbow tracks correspond to sequences shown in Movie S1 of the supplementary materials.

**Figure 4.**
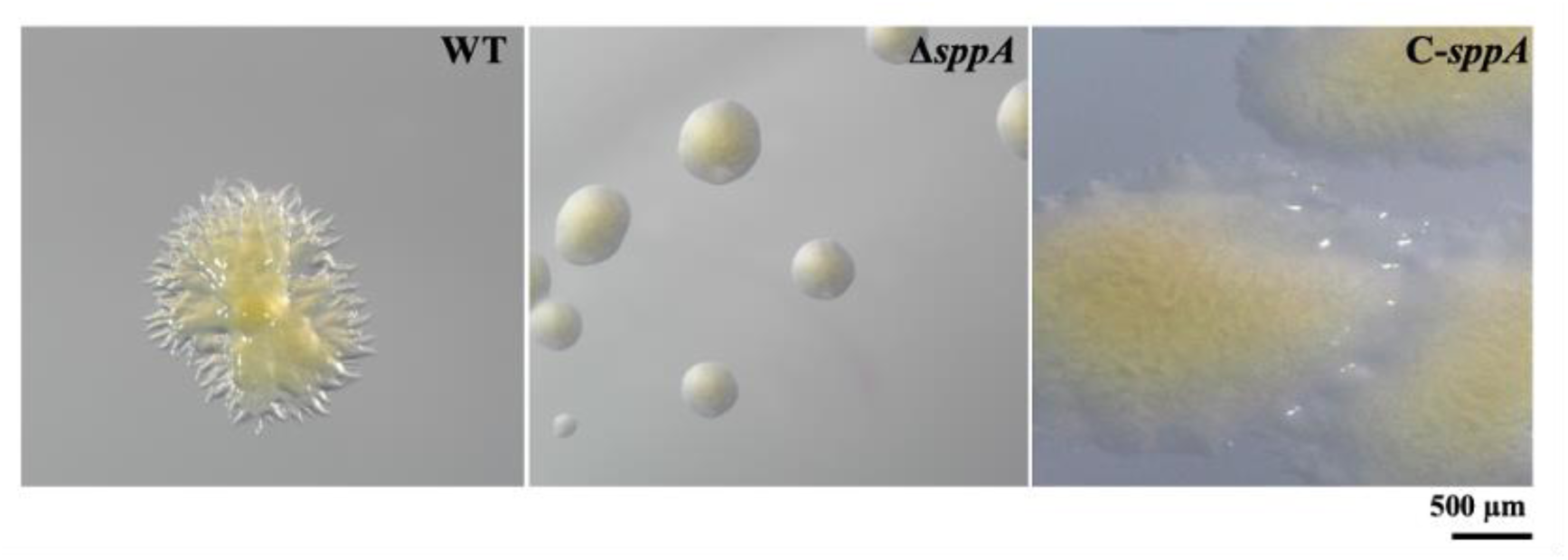
Photomicrographs of wild-type, mutant, and complemented *F. columnare* strains on MS agar. Colonies were cultured for 48 h at 28°C on MS agar. Strains examined were wild-type (WT), Δ*sppA,* and C-*sppA*. Photomicrographs were taken with a Nikon SMZ25 microscope. Images for WT and C-*sppA* strains were characterized for spreading colonies on MS agar, whereas the image of the Δ*sppA* mutant formed non-spreading colonies.

### The Δ*sppA* mutant exhibits defects in adhesion and biofilm formation

Given that gliding motility facilitates initial surface contact and colonization, we next investigated whether the impaired motility of the *sppA* mutant would affect its ability to adhere and form biofilms. Adhesion assays revealed that the Δ*sppA* mutant was significantly impaired in its ability to attach to polystyrene surfaces relative to the wild-type strain (Fig. 5A). Adhesion of cells to a surface is the first step in biofilm formation, and *F. columnare* forms biofilms on various surfaces including polystyrene. Crystal violet staining revealed that the Δ*sppA* mutant was partially deficient in biofilm formation compared to the wild-type strain, while the complemented strain showed intermediate biofilm biomass between the mutant and wild-type levels (Fig. 5B).

**Figure 5.**
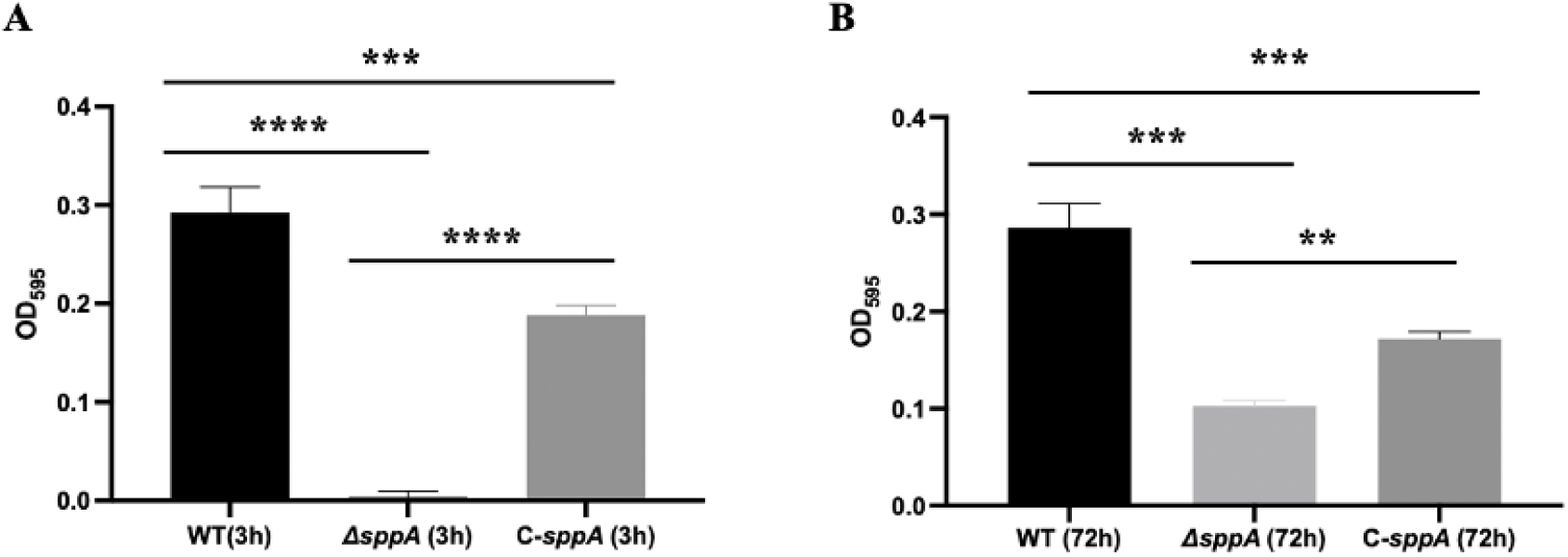
Adhesion and biofilm formation of wild-type *F. columnare* (WT), Δ*sppA* mutant, and Δ*sppA* mutant complemented with pCP23-SPPA. **(A)** Adhesion to polystyrene after 3 h of incubation at 28°C without shaking as determined by staining with crystal violet and measuring absorbance at 595 nm. The results are presented as mean ± SD, with each condition tested in quadruplicate (n = 4). Significant differences between groups were determined using one-way ANOVA, where \*\*\**P < 0.001* and \*\*\*\**P < 0.0001* indicate levels of statistical significance. **(B)** Biofilm formation on polystyrene of cells in MS broth incubated for 72 h at 28°C without shaking. The results are presented as mean ± SD, with each condition tested in quadruplicate (n = 4). Significant differences between groups were determined using one-way ANOVA, where \*\**P < 0.01* and \*\*\**P < 0.001* indicate levels of statistical significance.

### Enhanced outer membrane vehicles (OMVs) production in the Δ*sppA* mutant

To assess whether the loss of *sppA* affects cell envelope integrity and protein secretion, we analyzed the cellular ultrastructure using TEM. Significant ultrastructural differences between the Δ*sppA* mutant strain and the wild-type *F. columnare* were observed. The mutant strain showed a significant increase in the number of OMVs around the cells and a clearer cell membrane boundary (Fig. 6A). Quantitative analysis further confirmed that the number of OMVs per cell of the Δ*sppA* mutant strain (5.261 ± 1.512) was significantly increased by about 3.8-fold compared with that of the wild type (1.383 ± 0.5439) (\*\*\*\**P < 0.0001*) (Fig. 6B). These results suggest that deletion of the *sppA* gene may lead to the accumulation of intracellular protein products, which in turn expel excess or misfolded proteins by increasing the production of OMVs, as well as triggering the remodeling of the cell membrane structure.

**Figure 6.**
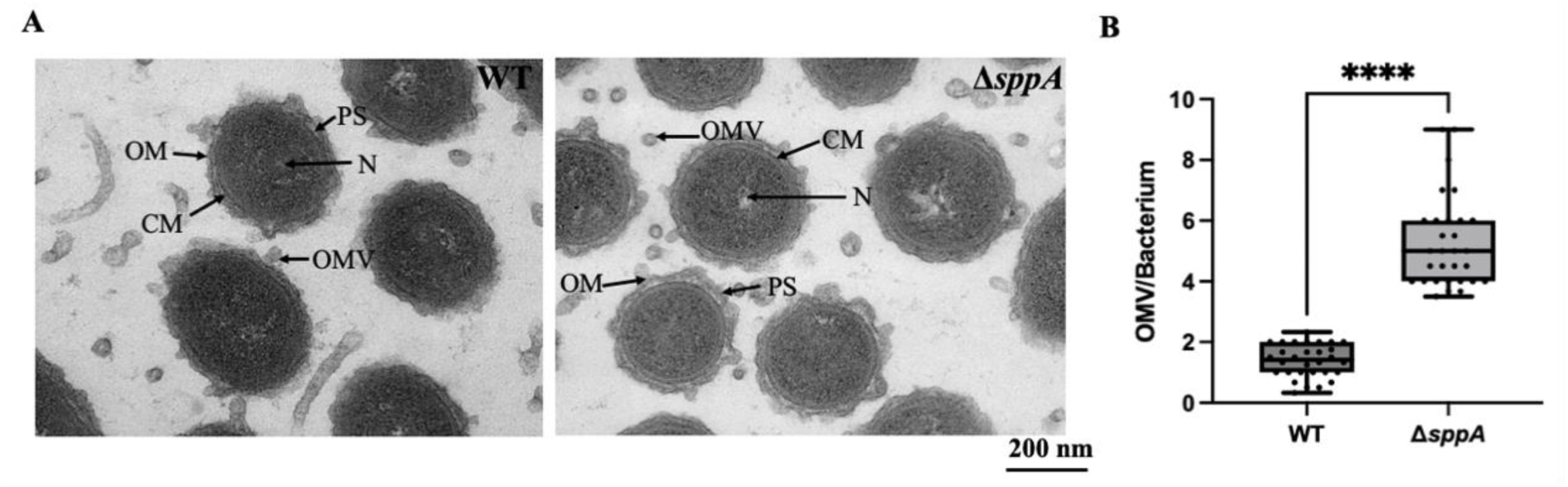
TEM micrographs of wild-type *F. columnare* and the *sppA* deletion mutant. **(A)** TEM observations of wild-type *F. columnare* and *sppA* gene deletion mutant. Arrows indicate nucleoid (N), cell membrane (CM), outer membrane (OM), outer membrane vesicle (OMV), and periplasmic space (PS). Scale bars represent 200 nm. **(B)** Comparative analysis of OMVs production per cell between wild-type strain and Δ*sppA* mutant strains of *F. columnare*. Data was calculated based on examining 60 randomly selected images, and is presented as means ± SD. **** *P < 0.0001*.

### Virulence of the Δ*sppA* mutant was reduced in the freshwater Medaka infection model

To evaluate the role of *sppA* in pathogenesis, we compared the virulence of wild-type *F. columnare* and the Δ*sppA* mutant in freshwater Medaka fish (Fig. 7). The artificial infection experiment revealed that on day 11 post-infection, the survival rate in the Δ*sppA* mutant challenge group reached 74.0%, compared to only 54.3% in the wild-type strain treatment group, indicating a significantly higher survival rate in the *sppA* gene knockout group. Fish infected with the Δ*sppA* mutant developed similar columnaris disease symptoms with the wild type counterpart, including yellow external lesions and lethargic swimming, during the 11-day observation period. *F. columnare* colonies were successfully recovered onto tobramycin-containing MS plates from various organs of dead fish, which confirmed that mortality resulted from columnaris infection.

**Figure 7.**
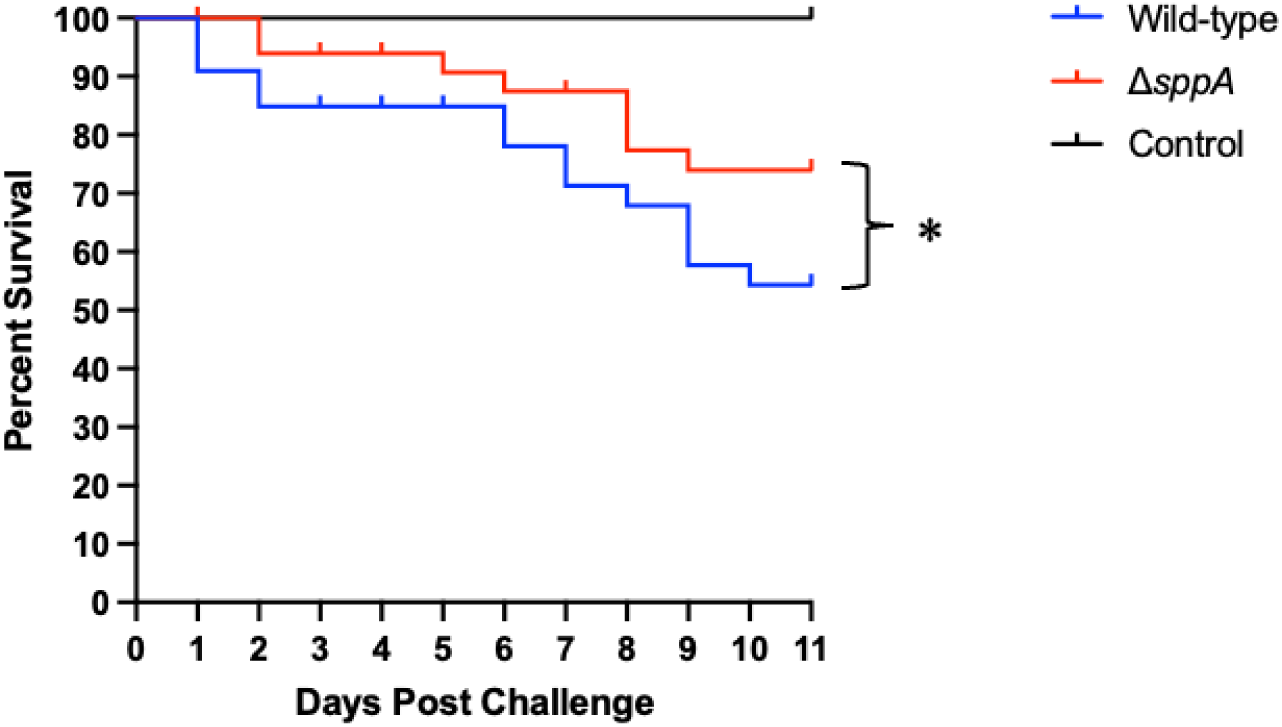
Virulence assessment of *F. columnare* wild type and Δ*sppA* mutant in Medaka fish. Medaka fish were subjected to exposure to the wild type (indicated in blue) and Δ*sppA* mutants (shown in red). A control group was exposed to an equivalent volume of MS growth medium (shown in black). Survival data was conducted using Kaplan-Meier log rank survival analysis, and the resulting survival curves were compared using an unpaired two-tailed Student’s t-test.

### Transcriptomic analysis revealed compensatory pathways for the peptide metabolites secretion

RNA-seq analysis identified 22 DEGs between wild-type and the Δ*sppA* mutant; *sppA* (the deleted locus) was the sole downregulated gene, and all remaining DEGs (n = 21) were upregulated (Table 3 and Fig. 8AB). Notably, genes encoding the MacA/MacB/TolC tripartite efflux pump (e.g., *macA*, *macB*, *tolC*) were significantly upregulated in the Δ*sppA* mutant. This RND-family transporter system is a complex protein secretion system found in bacteria that helps expel toxic compounds, including antibiotics, out of the cell. Since *sppA* functions as a protease involved in protein processing and degradation, its absence may lead to the accumulation of misfolded proteins or toxic metabolites, and the MacA/MacB/TolC tripartite efflux pump likely serves as a compensatory mechanism for enhanced efflux pump activity to manage cellular stress resulting from the *sppA* deficiency.

**Figure 8.**
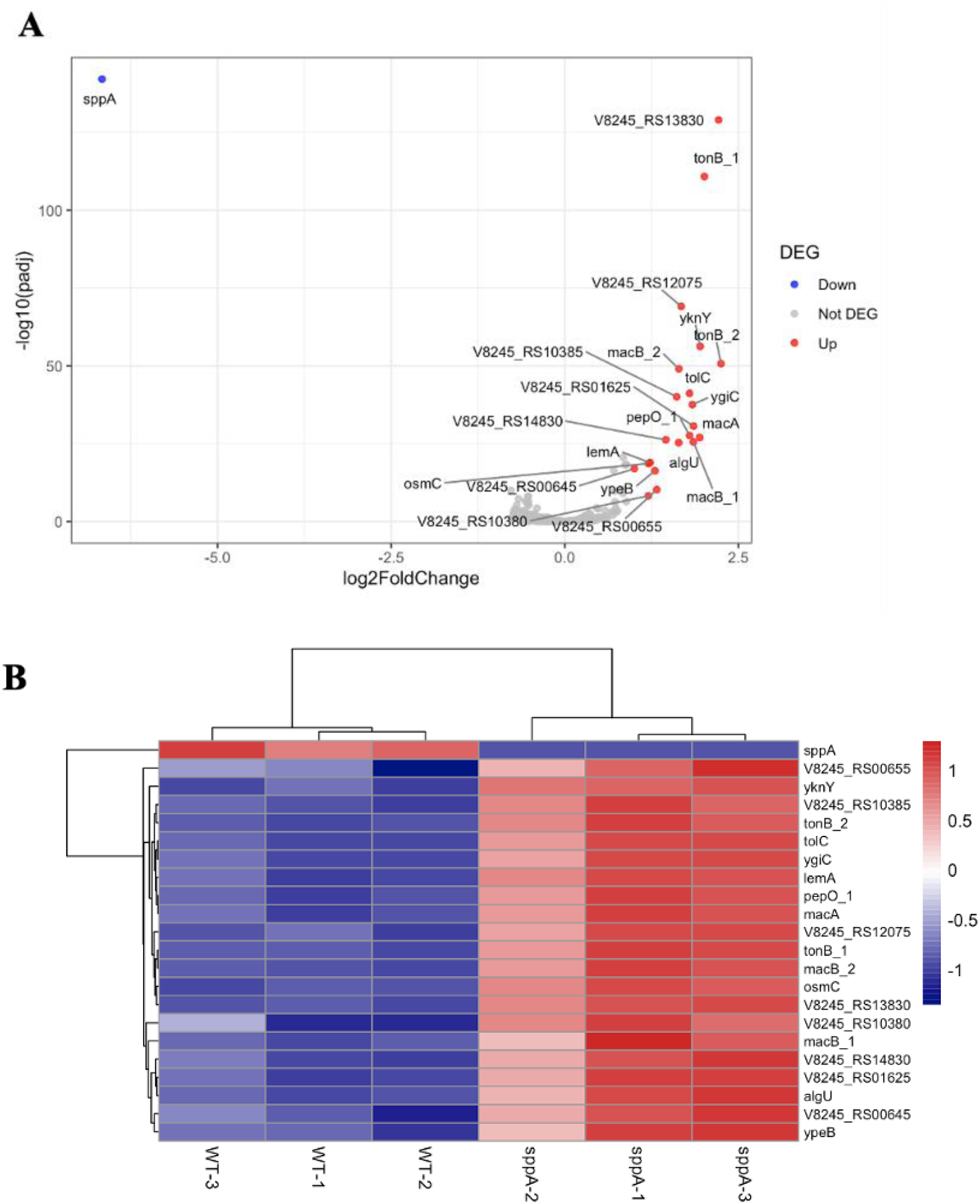
Statistics of differentially expressed genes between wild-type *F. columnare* and Δ*sppA*. **(A)** Volcano plot of DEGs comparing Δ*sppA* vs. the WT, p_adj < 0.05 and|log2FoldChange|>1. The x-axis represents the log2FoldChange, while the y-axis represents the statistical significance of each gene. **(B)** Heatmap of 22 DEGs between wild-type and Δ*sppA* groups. Each row represents a gene, and each column represents a sample, including control (WT-1, WT-2, WT-3) and treatment (*sppA*-1, *sppA*-2, *sppA*-3) groups. Gene expression levels were standardized using z-score normalization, with red indicating high expression and blue indicating low expression.

The upregulation of *algU*, encoding an extracytoplasmic function sigma factor, provides further evidence of stress response activation in the Δ*sppA* mutant. In many bacteria, AlgU-type sigma factors (*algU*) are known to stimulate OMV production and facilitate the removal of periplasmic proteins. This finding correlates with our TEM observations showing increased OMV formation in the Δ*sppA* mutant, suggesting that the OMVs may serve as an adaptive response to manage the cellular stress caused by *sppA* deficiency. Additionally, RNA-seq data revealed differential expression of genes involved in membrane homeostasis (*tolC*, *macA*, *macB*, *macB_2*, *lemA*, *tonB*), protein processing and degradation (*pepO_1*), and oxidative stress response (*algU*, *osmC*, *ypeB*, *yknY*, TPM domain-containing gene), indicating that *sppA* plays a broader role in maintaining cellular integrity via maintaining its proteolytic function.

Based on the experimental findings and the illustrated model, the deletion of *sppA* in *F. columnare* triggers a cascade of cellular responses to cope with signal peptide accumulation. Following *sppA* knockout, signal peptides that would normally be degraded accumulate in the inner membrane, creating membrane stress and potentially affecting protein translocation efficiency. This accumulation appears to induce compensatory mechanisms, as evidenced by the significant upregulation of the MacAB-TolC efflux pump system, which may serve as an alternative route for removing accumulated peptides or toxic compounds resulting from membrane stress. The increased production of OMVs, confirmed by TEM observations, likely represents another stress response mechanism to alleviate membrane perturbation and potentially export misfolded or aggregated proteins. The activation of the *algU* stress response pathway further confirms that *sppA* deletion creates significant cellular stress. Collectively, these findings demonstrate that *sppA* plays a central role in maintaining cellular homeostasis, and its absence forces the cell to activate multiple stress response pathways, particularly enhanced efflux pump activity and OMV production, to cope with membrane stress (Fig. 9).

**Figure 9.**
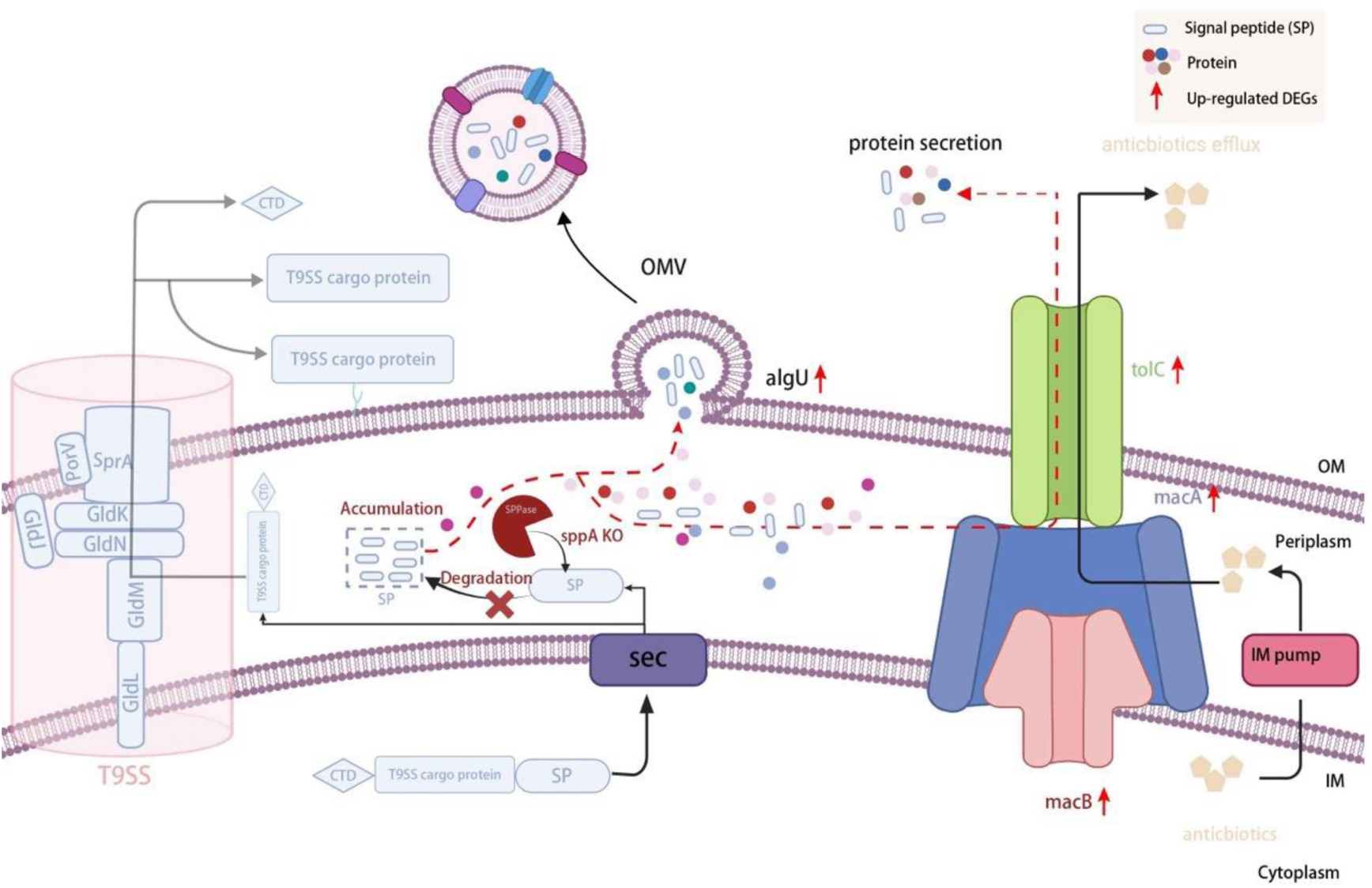
Proposed diagram for protein secretion and metabolites efflux in the periplasmic space of Δ*sppA* mutant. IM, inner membrane; OM, outer membrane; SP, signal peptide; SPPase, signal peptide peptidase; OMV, outer membrane vesicle; T9SS, type IX secretion system.

## Discussion

*F. columnare* represents a major bacterial pathogen affecting commercially important fish species, such as farmed catfish and tilapia, with substantial economic impacts on global freshwater aquaculture [17–21]. However, its virulence factors have been largely unexplored. Signal peptide peptidases play an essential role in protein processing by degrading N-terminal signal peptides that have been cleaved from newly synthesized proteins. This degradation is a critical step for maintaining proper protein translocation and localization by preventing the accumulation of signal peptide fragments. The activity of signal peptide peptidases is potentially important for the function of the T9SS in *F. columnare*, which mediates the export of multiple virulence factors, including proteases and adhesins [6]. The maturation and biological activity of these secreted virulence proteins depend on signal peptidase-mediated processing [22]. In the absence of *sppA*, signal peptidase activity would also be negatively affected due to the accumulation of signal peptides [15], which could potentially have an impact on bacterial pathogenicity. In this study, we successfully constructed a mutant strain (Δ*sppA*) with stable genetic characteristics using in-frame deletion of the gene *sppA* (encoding signal peptide peptidase A) by homologous recombination. This genetic construct enabled a systematic investigation of the enzyme’s role in bacterial pathogenesis and physiological functions, particularly its contribution to virulence factor processing. Using multiple analytical approaches, including ultrastructural characterization, growth kinetics, biofilm quantification, and global transcript profiling, our results revealed a pleiotropic regulatory role of *sppA* in *F. columnare*, with mutant leading to significant physiological alterations, such as impaired growth kinetics, attenuated biofilm development, enhanced outer membrane vesiculation, reduced virulence and activation of enhanced efflux pump activity to compensate for the *sppA* deficiency.

Transcriptome analysis showed that deletion of the *F. columnaris sppA* gene significantly altered the expression pattern of membrane-associated genes, confirming the key regulatory role of this gene in maintaining bacterial membrane homeostasis. In some bacterial pathogens, *sppA* cleaves signaling peptides released or extracted from membranes, and *sppA* deficiency may affect the ability to degrade signaling peptides remaining in the cell membrane [23]. Accumulation of these signaling peptides in the periplasm of the cell may interfere with protein translocation and affect cell membrane integrity through the Sec machinery [24, 25]. For example, studies in *B. subtilis* have shown that *sppA* knockout strains have significantly reduced levels of target protein secretion, a result that directly confirms the important role of the *sppA* gene in the protein secretion process in this bacterium [15]. Our study revealed that the Δ*sppA* mutant strain exhibited significant upregulation of an ECF σ factor (log2FC=1.64) and the stress response protein OsmC (log2FC=1.20). In addition, TEM analysis revealed a 3.8-fold increase (*p<0.0001*) in OMV production, collectively indicating a severe stress response in cells. These findings were consistent with the mechanism reported by MacDonald and Kuehn [26], where the *algU* σ factor alleviates bacterial stress by promoting OMV formation, demonstrating a conserved membrane stress response strategy among Gram-negative bacteria. Previous report suggested that OMV biogenesis serves not only as a passive consequence of membrane stress, but rather as an active mechanism for eliminating accumulated misfolded proteins [27]. Under the proteotoxic stress caused by *sppA* deletion, bacteria significantly enhance their survival by selectively packaging misfolded proteins into OMVs for extracellular disposal. Notably, *osmC* upregulation may assist in protein refolding under membrane stress conditions [28], promoting clearance of accumulated periplasmic proteins. Furthermore, up-regulation of the MacAB-TolC efflux pump may help bacteria excrete signal peptide fragments or other toxic molecules, thereby relieving pressure on the membrane system [29].

The investigation into the effects of the *sppA* gene deletion on *F. columnare* provides significant insights into its role in virulence and biofilm formation, which are important for understanding and managing columnaris disease in aquaculture settings. Our findings indicate that the deletion of the *sppA* gene significantly reduces the virulence of *F. columnare* in freshwater Medaka fish under the tested conditions. This could possibly be due to the disrupted protein system and elevated membrane stress in the knockout mutant. Previous study found that σE/AlgU activation inhibited the expression of virulence genes as a protective response [30]. This trade-off causes bacteria to prioritise maintaining cellular homeostasis over virulence factor production under membrane stress conditions, resulting in a partial reduction in virulence in freshwater Medaka. This result demonstrates that though the attenuation is moderate, *sppA* contributes to *F. columnare* pathogenicity and functions as a necessary virulence factor.

The Δ*sppA* mutant strain exhibited a significant increase in OMV production and a noticeable reduction in biofilm formation, which was highly consistent with the changes in expression of several membrane-associated genes in the transcriptomic data. Notably, the up-regulation of two TonB-dependent receptor genes and the cell morphology-determining protein gene suggested adjustments in envelope functions. These findings support the hypothesis that *sppA* maintains envelope integrity and protein homeostasis to sustain surface-associated functions. The investigation into the effects of the *sppA* gene deletion on *F. columnare* provides significant insights into its role in virulence and biofilm formation, which are important for understanding and managing columnaris disease in aquaculture settings.

In conclusion, our findings provide new insights into the biological roles of *sppA* in *F. columnare*. Our findings reveal that *sppA* is essential for maintaining membrane homeostasis and normal cellular physiology; the partial reduction in virulence indicates that *sppA* functions as an important virulence factor in *F. columnare* pathogenesis. These results provide important insights into the biological function of *sppA* in *F. columnare* and highlight the complex relationship between bacterial protein secretion, membrane integrity, and pathogenesis.

## Materials and methods

### Bacterial strains, plasmids, and growth conditions

*F. columnare* strain HLCZX-1 isolated from diseased mandarin fish (*Siniperca chuatsi*) was utilized as the wild-type strain for genetic analysis [31]. This strain was previously classified as genetic group 1 within the *F. columnare* species [32], and has been identified as *F. columnare* according to the new classification system [33]. *F. columnare* strains were cultured in modified Shieh (MS) agar or broth with shaking at 125 rpm at 28°C for 24 - 48 h [21]. *E. coli* strains DH5α and S17-1 λ *pir*, used for plasmid construction and bacterial conjugation, respectively, were grown at 37℃ and 250 rpm in lysogeny broth (LB) [34]. Plasmids in *E. coli* were selected using 100 μg/mL ampicillin, while selection for plasmids in *F. columnare* was achieved with 5 μg/mL tetracycline. In addition, 1 μg/mL of tobramycin was employed to counterselect against *E. coli* for conjugation experiments. The experimental strains and plasmids used in this study are listed in Table 1, and primers are listed in Table 2.

**Table 1.**
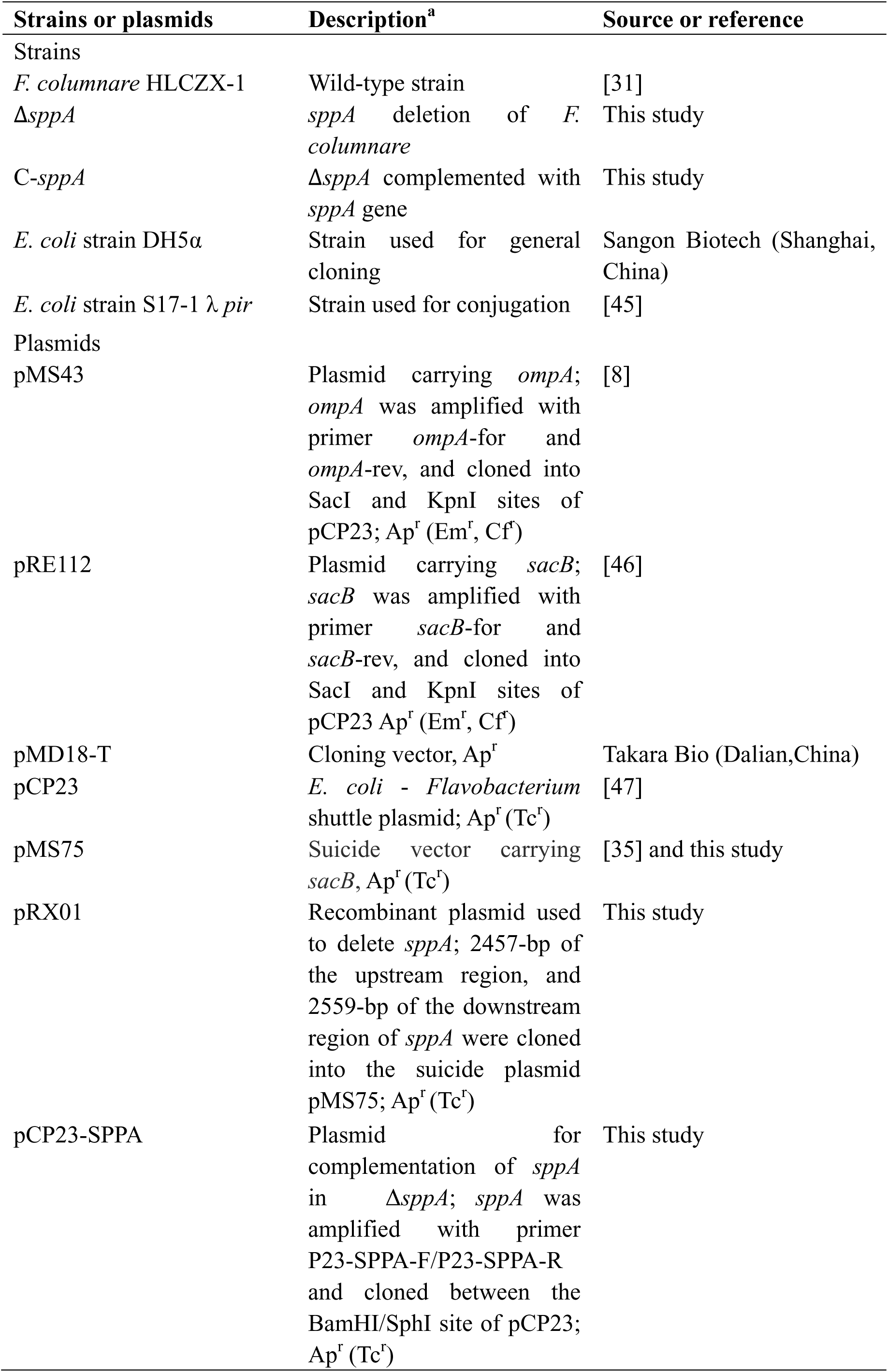
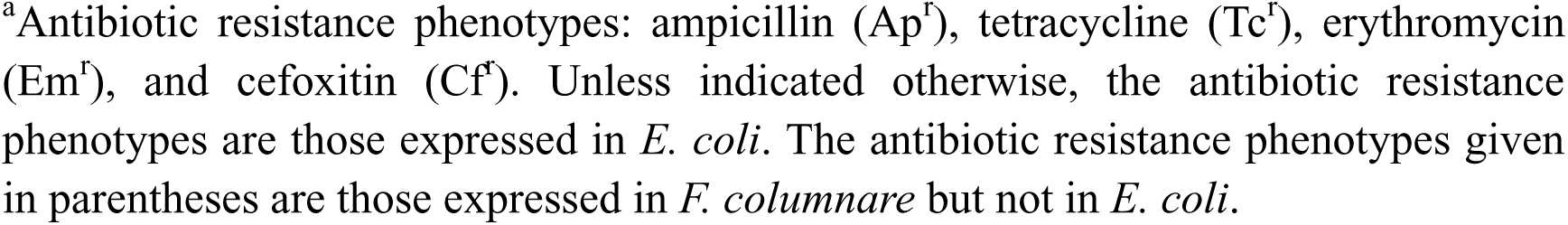
Bacterial strains and plasmids used in this study.

| Strains or plasmids | Description<sup>a</sup> | Source or reference |
| --- | --- | --- |
| **Strains** |  |  |
| *F. columnare* HLCZX-1 | Wild-type strain | [31] |
| $\Delta sppA$ | *sppA* deletion of *F. columnare* | This study |
| C-*sppA* | $\Delta sppA$ complemented with *sppA* gene | This study |
| *E. coli* strain DH5α | Strain used for general cloning | Sangon Biotech (Shanghai, China) |
| *E. coli* strain S17-1 $\lambda$ pir | Strain used for conjugation | [45] |
| **Plasmids** |  |  |
| pMS43 | Plasmid carrying *ompA*; *ompA* was amplified with primer *ompA*-for and *ompA*-rev, and cloned into SacI and KpnI sites of pCP23; Ap<sup>r</sup> (Em<sup>r</sup>, Cf<sup>r</sup>) | [8] |
| pRE112 | Plasmid carrying *sacB*; *sacB* was amplified with primer *sacB*-for and *sacB*-rev, and cloned into SacI and KpnI sites of pCP23 Ap<sup>r</sup> (Em<sup>r</sup>, Cf<sup>r</sup>) | [46] |
| pMD18-T | Cloning vector, Ap<sup>r</sup> | Takara Bio (Dalian, China) |
| pCP23 | *E. coli* - *Flavobacterium* shuttle plasmid; Ap<sup>r</sup> (Tc<sup>r</sup>) | [47] |
| pMS75 | Suicide vector carrying *sacB*, Ap<sup>r</sup> (Tc<sup>r</sup>) | [35] and this study |
| pRX01 | Recombinant plasmid used to delete *sppA*; 2457-bp of the upstream region, and 2559-bp of the downstream region of *sppA* were cloned into the suicide plasmid pMS75; Ap<sup>r</sup> (Tc<sup>r</sup>) | This study |
| pCP23-SPPA | Plasmid for complementation of *sppA* in $\Delta sppA$; *sppA* was amplified with primer P23-SPPA-F/P23-SPPA-R and cloned between the BamHI/SphI site of pCP23; Ap<sup>r</sup> (Tc<sup>r</sup>) | This study |

<sup>a</sup>Antibiotic resistance phenotypes: ampicillin (Ap<sup>r</sup>), tetracycline (Tc<sup>r</sup>), erythromycin (Em<sup>r</sup>), and ceftiofur (Cf<sup>r</sup>). Unless indicated otherwise, the antibiotic resistance phenotypes are those expressed in *E. coli*. The antibiotic resistance phenotypes given in parentheses are those expressed in *F. columnare* but not in *E. coli*.

**Table 2.**
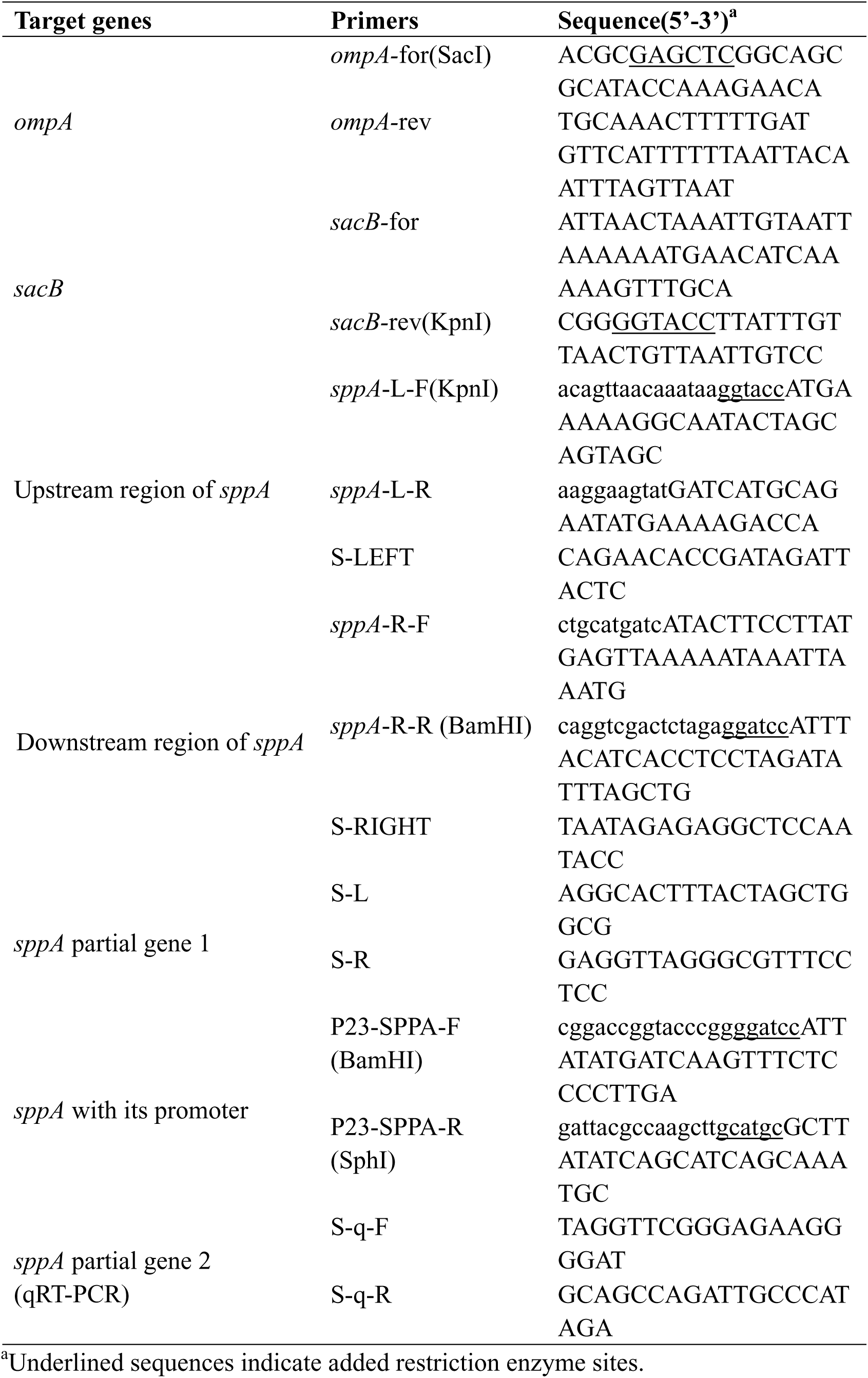
Sequences of primers used in this research.

**Table 3.** Representative genes differentially expressed in the *sppA* mutant compared with the wild-type strain.

| Gene symbol | Function description | log2FC<br>(Δ*sppA*/WT) |
| --- | --- | --- |
| **Resistance** |  |  |
| *algU* | RNA polymerase ECF-type sigma factor | 1.64 |
| *V8245_RS12075* | Beta-carotene 15,15'-monooxygenase | 1.68 |
| *osmC* | putative stress-induced protein OsmC | 1.20 |
| *ypeB* | DUF3820 domain-containing protein | 1.30 |
| *yknY* | ABC-type antimicrobial peptide transport system, ATPase component | 1.95 |
| *V8245_RS13830* | TPM domain-containing protein | 2.21 |
| **Cell membrane** |  |  |
| *tolC* | outer membrane efflux protein | 1.80 |
| *macA* | ABC transporter, RND-adaptor-like protein | 1.94 |
| *macB_1* | ABC-type antimicrobial peptide transport system, permease component | 1.85 |
| *macB_2* | Macrolide export ATP-binding/permease protein MacB | 1.64 |
| *V8245_RS00645* | Peptidase, M16 family protein | 1.00 |
| *lemA* | LemA family protein | 1.22 |
| *V8245_RS00655* | hypothetical protein | 1.32 |
| *V8245_RS10380* | Lipoprotein | 1.20 |
| *V8245_RS10385* | Lipoprotein | 1.61 |
| *ygiC* | Putative acid--amine ligase YgiC | 1.84 |
| *tonB_1* | Putative TonB-dependent receptor | 2.01 |
| *tonB_2* | Putative TonB-dependent receptor | 2.25 |
| *V8245_RS14830* | Glycosyl transferase, group 2 family | 1.45 |
| *V8245_RS01625* | Cell shape-determining protein | 1.85 |
| **Protein processing and degradation** |  |  |
| *pepO_1* | Neutral endopeptidase | 1.80 |
| *sppA* | Signal peptide peptidase A | -6.67 |

### Construction of pMS75 suicide plasmid for gene editing

Suicide vector pMS75 was re-constructed as previously described [35]. In brief, the promoter region of the *Flavobacterium johnsoniae ompA* gene was amplified by PCR using pAS43 as the template, with r-Taq DNA polymerase (Takara) and the primers *ompA*-for (introducing a SacI site) and *ompA*-rev. Similarly, the *sacB* gene was amplified from pRE112 using the primers *sacB-*for and *sacB-*rev (introducing a KpnI site). These two fragments were linked using overlap extension PCR and cloned them into the pMD-18T vector (Takara Bio, Dalian, China), adding the *ompA* promoter in front of the *sacB* gene to replace its original promoter. The purified PompA + *sacB* fragment, released from the pMD-18T vector after digestion with SacI and KpnI and proliferated in *E. coli* DH5α. The pMD-18T_sacB fragment was cut by the KpnI and BanHI and then ligated into the pCP23 vector, which had been digested with the same enzymes, to construct the suicide plasmid pMS75.

### Conjugative transfer of plasmids in *F. columnare* strains

Plasmids were transferred from *E. coli* S17-1 λ *pir* to *F. columnare* by conjugation as described previously [36]. Briefly, 50 µL of overnight *E. coli* culture was inoculated into 5 mL of LB broth, and 50 µL of overnight *F. columnare* culture was inoculated into 5 mL of MS broth. The cultures were incubated with shaking at 37°C and 28°C, respectively, until their optical densities at 600 nm (OD_600_) reached 0.5. The bacterial solution was then centrifuged at 8,000 × g for 1 min, followed by washing and resuspending in MS broth. Suspensions of *E. coli* and *F. columnare* were mixed and then centrifuged again at 8,000 × g for 1 min to remove excess medium. The cell mixture was resuspended in 200 µL of MS broth, spotted on MS agar, and incubated at 30°C for 24 h. After incubation, the cells were scraped off from the agar and resuspended in 1 mL of MS broth. Then 100-µL aliquots were spread on MS agar containing 1 µg/mL tobramycin and 5 µg/mL tetracycline and incubated at 28°C for 72 h.

### Construction of *sppA* deletion mutant

In-frame deletion was generated using established protocols [35]. Briefly, a 2457 bp of the upstream fragment of *sppA* was amplified from *F. columnare* genomic DNA using the primer pair *sppA*-L-F (containing the KpnI site) and *sppA*-L-R, while a 2559 bp downstream fragment was amplified using *sppA*-R-F and *sppA*-R-R (containing the BamHI site) (Table 2). The amplified product was assembled with linearized pMS75 that was digested with KpnI and BamHI using ClonExpress Ultra One Step Cloning Kit V2 (Vazyme Biotech, Nanjing), resulting in a recombinant plasmid pRX01. The plasmid pRX01 was introduced into the *F. columnare* wild-type strain via conjugation. Colonies with the plasmid integrated into the cell were selected based on tetracycline resistance. These colonies were streaked onto MS agar with tetracycline, and isolated colonies were subsequently cultured in liquid MS medium without tetracycline to facilitate plasmid loss by DNA recombination. The cultures were then plated on MS medium containing 10% sucrose, allowing colonies that had lost the *sacB*-containing plasmid to grow, since *sacB* is lethal to cells in the presence of sucrose. Finally, the *sppA* deletion mutant was verified in the colonies growing with sucrose by PCR, and the mutant was designated as Δ*sppA*.

### Complementation of the *sppA* mutant

A 2229 bp fragment spanning the *sppA* gene was amplified using the primers P23-SPPA-F (introducing the BamHI site) and P23-SPPA-R (introducing the SphI site) (Table 2). The amplified product was ligated into the shuttle vector pCP23 to produce a recombinant plasmid pCP23-SPPA, followed by introduced into the *F. columnare sppA* mutant via conjugation.

To further confirm the presence of the *sppA* gene, its expression level was quantified using quantitative real-time PCR (qRT-PCR) on an Applied Biosystems QuantStudio 7 Pro system (Thermo Fisher) with gene-specific primers S-q-F and S-q-R (Table 2). Total RNA was extracted using the RNeasy Mini Kit (QIAGEN) and quantified by NanoDrop 2000 spectrophotometry (Thermo Fisher). cDNA synthesis was performed using HiScript IV All-in-One Ultra RT SuperMix (Vazyme) following the manufacturer’s protocol. qPCR reactions were carried out with Taq Pro Universal SYBR qPCR Master Mix (Vazyme) under the following cycling conditions: initial denaturation at 95°C for 30 s, followed by 40 cycles of 95°C for 10 s and 60°C for 30 s, with a final melt curve analysis of 95°C for 15 s, 60°C for 60 s and 95°C for 15 s. The gene *glyA* was used as an internal control, and relative gene expression was calculated using the 2^−ΔΔCT^ method [35].

### Growth curve analysis

*F. columnare* strains were revived from frozen stock by plating on MS agar and incubating at 28°C for 36 hours. The resulting colonies were used to inoculate 10 mL of MS medium, which was incubated overnight at 28°C with shaking at 125 rpm. For strains carrying plasmids, 2.5 μg/mL tetracycline was added to the medium. Overnight cultures of the wild type, *ΔsppA* mutant, and the complementary strains were adjusted to an OD₆₀₀ of 0.5 and used to inoculate 3 mL of fresh MS broth. The cultures were incubated at 28°C with shaking at 125 rpm, and OD₆₀₀ values were measured every 9 h for 54 h using a DEN-600 spectrophotometer (Biosan, Latvia).

### Antibiotic resistance test

The antimicrobial susceptibility of *F. columnare* wild-type and Δ*sppA* mutant strains was evaluated against three aquaculture-relevant antibiotics: oxytetracycline dihydrate (OTC), enrofloxacin (ENRO), and florfenicol (FF), which were obtained from Aladdin (Shanghai, China). The broth microdilution method was employed to determine minimum inhibitory concentrations (MICs) following standardized procedures. Overnight bacterial cultures were adjusted to an OD_600_ of 0.04 ± 0.005 in MS broth before testing. Two-fold serial dilutions of each antibiotic were prepared in 96-well microtiter plates (Corning 167008, Corning, NY), covering a concentration range from 16 μg/mL to 0.00375 μg/mL. Each well received 100 μL of antibiotic solution, followed by inoculation with 20 μL of bacterial suspension, with all concentrations tested in triplicate. Following a 48-hour incubation at 28°C in the dark to prevent antibiotic degradation, MIC values were determined as the lowest antibiotic concentrations that completely inhibited visible bacterial growth.

### Adhesion and biofilm formation assay

Bacterial adhesion to polystyrene was quantified using a crystal violet staining assay as described previously [36]. *F. columnare* cultures were grown in MS broth to an OD_600_ of 0.5. Cells were harvested by centrifugation (1 mL per sample), washed, and resuspended in sterile distilled water. Cell suspensions (100 µL) were added to wells of a 96-well flat-bottom polystyrene plate (Corning 167008, Corning, NY) and incubated statically at 28°C for 3 h. After incubation, non-adherent cells were removed by washing twice with sterile distilled water. Adherent cells were stained with 100 µL of 1% (w/v) crystal violet for 30 min at room temperature, followed by four washes with sterile distilled water. The bound dye was solubilized in 100 µL of absolute ethanol, and absorbance was measured at 595 nm using a SpectraMax iD3 Multi-Mode Microplate Reader (Molecular Devices, USA).

The biofilm formation capability in wild-type, mutant, and complemented strains was assessed using the methodologies described in previous research [37]. In brief, cells were grown in MS medium to the mid-logarithmic phase (OD₆₀₀ = 0.5). The cultures were diluted 1:100 in MS medium, and 150 μL of the diluted culture was added to each well of a 96-well flat-bottom polystyrene microplate (Corning 167008, Corning, NY). The plate was covered with aluminum foil and incubated at 28°C for 72 h. Biofilm formation was tested in four wells per strain, with sterile, uninoculated medium serving as a negative control. After incubation, the medium was discarded, and the wells were washed three times with 200 μL of sterile distilled water. Each well was then stained with 150 μL of 1% (w/v) crystal violet solution for 30 minutes at room temperature. Excess dye was removed by washing the wells four times with 200 μL of sterile distilled water. The bound crystal violet was dissolved with 100 μL of ethanol, and absorbance was measured at 595 nm (OD₅₉₅) using a SpectraMax iD3 Multi-Mode Microplate Reader (Molecular Devices, USA). The absorbance of the uninoculated negative control was subtracted from the absorbance of each strain.

### Analysis of cell motility and colony morphology

Gliding of individual cells was observed microscopically after cells were grown with shaking at 28°C in 1/10 MS broth overnight with shaking; tunnel slides were prepared using double-sided tape, glass microscope slides, and coverslips as previously described [38], then 10 μl of culture was added to each tunnel, incubated for 5 min, and cell motility was recorded at 25°C using a Nikon Ci-L plus microscope equipped with an SC2000C CMOS camera, with rainbow traces of cell movement generated using Fiji version 2.14.0 (https://imagej.net/software/fiji/) and the Color FootPrint macro [39]. Colony morphology was examined by serial dilution plating. In brief, mid-exponential phase cultures were serially diluted (10-fold dilutions) in sterile MS medium, and 100 µL of the appropriate dilutions were spread onto MS agar plates. After incubation at 28°C for 48 h, individual colony morphology was observed and photographed using a Nikon SMZ25 microscope (Tokyo, Japan).

### Transmission electron microscopy (TEM) analysis

For TEM analysis, bacterial cells from triplicate overnight cultures were harvested by centrifugation and immediately fixed in TEM-grade fixative at 4°C. The fixed cells were washed three times with 0.1 M phosphate buffer (PB, pH 7.4) for 3 minutes per wash, resuspended in 1% agarose solution, and encapsulated before the agarose solidified. The agarose blocks were post-fixed with 1% osmium tetroxide (OsO₄) in 0.1 M PB (pH 7.4) for 2 hours at room temperature in the dark, followed by three rinses in 0.1 M PB. The samples were dehydrated at room temperature through a graded ethanol series (30%, 50%, 70%, 80%, 95%, and 100% ethanol) and two changes of acetone. Resin infiltration was performed with acetone and EMBed 812 resin mixtures (1:1 for 2-4 hours, 1:2 overnight, and pure resin for 5-8 hours) at 37°C, after which the samples were embedded in pure resin and cured overnight at 37°C. Polymerization was carried out at 60°C for over 48 hours. Ultrathin sections (60-80 nm) were prepared using an ultramicrotome, then placed on 150-mesh copper grids coated with formvar film and stained with 2% uranyl acetate and 2.6% lead citrate. The stained sections were observed under TEM, and detailed images were captured.

### Fish challenge

Wild-type *F. columnare* and Δ*sppA* mutant strains were cultured overnight in MS medium for 24 h at 28°C. On the following day, 50 μl of these cultures were transferred into 5 ml of fresh MS broth and incubated with shaking at 28°C until the optical density at 600 nm (OD_600_) reached 0.4. To quantify viable cells, serial dilutions (in triplicate) of the cultures were plated and enumerated on MS agar. Plates were incubated for 48 h at 28°C.

Freshwater medaka fish were hatched and obtained from the fish husbandry unit of the State Key Lab of Moraine Pollution (City University of Hong Kong) and were used for bacterial challenge. No signs of disease were observed before the challenge, and no evidence of *F. columnare* infection was detected in the uninfected control tanks and acclimation tanks throughout the study. Fish were transferred to challenge aquaria one week prior to the immersion challenge for acclimation.

Challenges were performed using triplicate 1-liter beakers with restricted water flow at 28°C, each containing 10 fish. For the immersion challenge, water flow was stopped, and bacterial cultures were added to the beakers, which were then incubated for 2 h before resuming water flow. Control beakers were inoculated with MS broth instead of bacterial cultures. The final challenge concentrations for the experiment were 7×10^5^ CFU/mL for the wild-type *F. columnare* and 1.05 × 10^6^ CFU/mL for the Δ*sppA* mutant. Mortalities were monitored, removed, and recorded daily. Data from the triplicate beakers for each strain were combined, and survivor fractions were calculated. To confirm the presence of *F. columnare*, the deceased fish were randomly selected and subjected to bacterial examination. Swabs from external and internal organs were streaked on MS agar to identify the plausible yellow, rhizoid, and adherent colonies of *F. columnare*. The present experiment was conducted in compliance with the animal research guidelines of the Hong Kong Special Administrative Region (HKSAR) under animal license [Ref No.: (24-50) in DH/HT&A/8/2/5 Pt.14] and with approval from the City University Animal Ethics Committee (Approval No.: A-0402).

### RNA isolation and transcriptomic analysis

Wild-type *F. columnare* and the Δ*sppA* mutant were cultured in MS broth until they reached early stationary phase (OD_600_ = 0.8). Total RNA was extracted using the RNeasy Mini Kit (QIAGEN, Valencia, CA) and subsequently treated with DNase I to eliminate any contaminating DNA. The quality and quantity of RNA were assessed using NanoDrop and Agilent 2100 instruments (Agilent Technologies, Palo Alto, CA). Three biological replicates were conducted for each strain. Ribosomal RNA depletion and library construction were conducted by the Novogene company (Beijing, China). Quantified libraries were pooled and sequenced on Illumina NovaSeq X Plus to generate the raw data of each sample. The qualities of the resulting raw reads were first evaluated by the FastQC program (v0.11.8) [40]. Low-quality sequences and adapters were trimmed using Trimmomatic v0.32 [41]. High-quality RNA-seq sequences were then mapped to the *F. columnare* reference genome (GCF_050711845.1) using Bowtie2 [42]. Gene expression levels were quantified using FeatureCounts software (v2.0.3) [43]. Differentially expressed genes (DEGs) between the wild-type and Δ*sppA* strains were identified using DESeq2 (version 1.34.0) [44]. Significantly differentially expressed genes were considered by parameters: |log2(fold change)| > 1 and false discovery rate (FDR) less than 0.05 [21]. Raw sequencing reads were deposited in the NCBI Sequence Read Archive (SRA) as part of BioProject PRJNA1321558.

### Statistical analyses

All experiments were conducted with at least three biological replicates. Data were presented as mean ± standard deviation. Statistical analyses were performed using SPSS 16.0. Significance was determined by Student’s t-test or one-way ANOVA followed by LSD test. *P < 0.05* was considered statistically significant. GraphPad Prism version 10 (GraphPad Software, LLC) was also used to compute statistical tests.

## Declaration of competing interest

The authors declare that they have no known competing financial interests or personal relationships that could have appeared to influence the work reported in this paper.

## Acknowledgement

This study was supported by the APRC-CityU New Research Initiatives/Infrastructure Support from Central (9610574, 7006064), and Early career scheme (project number 9048294) from the Hong Kong Research Grant Council.

## Author contributions

**Ruoxi Zhu:** Conceptualization, Formal analysis, Investigation, Methodology, Visualization, Writing–original draft; Writing–review and editing. **Liang Zhong:** Conceptualization, Investigation. **Yuying Xun:** Data curation, Investigation, Writing– review and editing. **Shucheng Zheng:** Conceptualization, Methodology. **Yongtao Zhu:** Conceptualization, Methodology, Supervision, Writing–review and editing. **Wenlong Cai:** Conceptualization, Funding acquisition, Methodology, Supervision, Writing–review and editing.

## Supplementary materials

**Table S1.** The minimum inhibitory concentrations of the wild-type strain and the *SppA* gene deletion mutant.

**Movie S1:** Bacterial gliding motility (WT, *ΔsppA mutant*, and complementary strains) at the single-cell level under time-lapse microscopy.

